# Combining electrophysiology with MRI enhances learning of surrogate-biomarkers

**DOI:** 10.1101/856336

**Authors:** Denis Alexander Engemann, Oleh Kozynets, David Sabbagh, Guillaume Lemaitre, Gaël Varoquaux, Franziskus Liem, Alexandre Gramfort

## Abstract

Electrophysiological methods, i.e., M/EEG provide unique views into brain health. Yet, when building predictive models from brain data, it is often unclear how electrophysiology should be combined with other neuroimaging methods. Information can be redundant, useful common representations of multimodal data may not be obvious and multimodal data collection can be medically contraindicated, which reduces applicability. Here, we propose a multimodal model to robustly combine MEG, MRI and fMRI for prediction. We focus on age prediction as surrogate biomarker in 674 subjects from the Cam-CAN. Strikingly, MEG, fMRI and MRI showed additive effects supporting distinct brain-behavior associations. Moreover, the contribution of MEG was best explained by source-topography of power spectra between 8 and 30 Hz. Finally, we demonstrate that the model maintains benefits of stacking when data is missing. The proposed framework hence enables multimodal learning for a wide range of biomarkers from diverse types of brain signals.

## Introduction

Non-invasive electrophysiology assumes a unique role in clinical neuroscience. Magneto- and electophencephalography (M/EEG) have an unparalleled capacity for capturing brain rhythms without penetrating the skull. EEG can be readily operated in a wide array of peculiar situations, such as surgery (Baker et al., 1975), flying an aircraft (Skov and Simons, 1965) or sleeping (Agnew Jr et al., 1966). Compared to EEG, MEG captures a more selective set of brain sources with greater spectral and spatial definition (Ahlfors et al., 2010; Hari et al., 2000). Yet, neither of them is optimal for isolating anatomical detail. Clinical practice in neurology and psychiatry therefore relies on additional neuroimaging modalities with enhanced spatial resolution such as magnetic resonance imaging (MRI), functional MRI (fMRI) or positron emission tomography (PET). Recently, machine learning has received significant interest in clinical neuroscience for its potential to predict from such heterogeneous multimodal brain data (Woo et al., 2017).Unfortunately, the effectiveness of machine learning in psychiatry and neurology is constrained by the lack of large high-quality datasets (Woo et al., 2017; Varoquaux, 2017; Bzdok and Yeo, 2017; Engemann et al., 2018) and comparably limited understanding about the data generating mechanisms (Jonas and Kording, 2017). This, potentially, limits the advantage of complex learning strategies proven successful in purely somatic problems (Esteva et al., 2017; Yoo et al., 2019; Ran et al., 2019).

In clinical neuroscience, prediction can therefore be pragmatically approached with classical machine learning algorithms (Dadi et al., 2019), expert-based feature engineering and increasing emphasis on surrogate tasks. Such tasks attempt to learn on abundant high-quality data an outcome that is not primarily interesting, to then exploit its correlation with the actual outcome of interest in small datasets. This can be seen as transfer learning problem (Pan and Yang, 2009) which, in its simplest form, is implemented by reusing predictions from a surrogate-marker model as predictors in the small dataset. Over the past years, predicting the age of a person from its brain data has crystalized as a surrogate-learning paradigm in neurology and psychiatry (Dosenbach et al., 2010). First results suggest that the prediction error of models trained to learn age from brain data of healthy populations provides clinically relevant information (Cole et al., 2018; Ronan et al., 2016; Cole et al., 2015) related to neurodegenerative anomalies, physical and cognitive decline (Kaufmann et al., 2019). For simplicity, this characteristic prediction error is often referred to as the brain age delta or Δ(Smith et al., 2019). Can learning of such a surrogate biomarker be enhanced by combining expert-features from M/EEG, fMRI and MRI?

Research on aging has suggested important neurological group-level differences between young and elderly people: Studies have found alterations in grey matter density and volume, cortical thickness and fMRI-based functional connectivity, potentially indexing brain atrophy (Kalpouzos et al., 2012) and decline-related compensatory strategies. Peak frequency and power drop in the alpha band (8-12Hz), assessed by EEG, has been linked to aging-related slowing of cognitive processes, such as the putative speed of attention (Clark et al., 2004; Babiloni et al., 2006). Increased anteriorization of beta band power (15-30Hz) has been associated with effortful compensatory mechanisms (Gola et al., 2013) in response to intensified levels of neural noise, i.e., decreased temporal autocorrelation of the EEG signal as revealed by flatter 1/f slopes (Voytek et al., 2015). Importantly, age-related variability in fMRI and EEG seems to be independent to a substantial degree (Kumral et al., 2019).

The challenge of predicting at the single-subject level from such heterogenous neuroimaging modalities governed by distinct data-generating mechanisms has been recently addressed with model-stacking techniques (Rahim et al.,2015; Karrer et al., 2019; Liem et al., 2017). Rahim et al. (2015) enhanced classification in Alzheimer’s disease by combined fMRI with PET prediction stacking (Wolpert, 1992), however, such that the stacked models reflected input data from modalities. Liem et al. (2017) have then applied this approach to age-prediction and found that combining anatomical MRI with fMRI significantly helped reduce errors while facilitating detection of cognitive impairment. This suggests that stacked prediction might also enable combining MRI with electrophysiology. Yet, this idea faces one important obstacle related to the clinical reality of data collection. It is often not practical to do multimodal assessments for all patients. Scanners may be overbooked, patients may not be in the condition to undergo MRI and acute demand in intensive care units may dominate priorities. Incomplete and missing data is therefore inevitable and has to be handled to unleash the full potential of multimodal predictive models.

To tackle this challenge, we developed a stacking model to predict age from electrophysiology and MRI including any case for which there was the opportunity to see at least one modality. We therefore call it opportunistic stacking model. For validation, we chose the currently largest public multimodal imaging and electrophysiology resource: The Cam-CAN dataset contains rich neuropsychological data, magnetic resonance imaging as well as non-invasive high-resolution electrophysiology in the form of magnetoencephalography (MEG) for more than 650 healthy subjects between 17 and 90 years (Shafto et al., 2014; Taylor et al., 2017). The choice of MEG has the advantage of improved spatial and frequency resolution. This should help identify robust and translatable electrophysiology markers potentially suitable for clinical EEG. Therefore, our study focuses on the following questions: 1) Can MRI-based prediction of age be enhanced with electrophysiology? 2) Do fMRI and MEG carry non-redundant clinically relevant information? 3) What are the most informative electrophysiological markers of aging? 4) Can potential advantages of multimodal learning be maintained in the presence of missing values?

## Results

### Opportunistic prediction-stacking approach

We begin by summarizing the proposed method. To build a model for predicting age from electrophysiology, functional MRI and anatomical MRI, we employed prediction-stacking (Wolpert, 1992). As in Liem et al. (2017), here, the stacked models referred to different input data instead of alternative models on the same data. We used ridge regression (Hoerl and Kennard, 1970) to linearly predict age from high-dimensional inputs of each modality. Linear predictions were based on distinct features from anatomical MRI, fMRI and MEG that have been commonly associated with aging. For MEG, we extracted evoked response latencies, alpha band peak frequency, 1/f slope topographies, source-level spectral power topographies and bivariate functional connectivity. For MRI we included cortical thickness, cortical surface area and subcortical volume. For fMRI we computed functional connectivity from the time-series. For detailed description of the features, see Table 3, section Feature extraction in materials and methods. To correct for the necessarily biased linear model, we then used non-linear random forest regressor with age predictions from the linear model as lower-dimensional input features.

Thereby, we made sure to use consistent cross-validation splits for all layers and automatically selected central tuning-parameters of the linear model and the random forest with nested cross-validation. Our stacked models handle missing values by treating missing value as data, provided there is an opportunity to see at least one modality (Josse et al., 2019). We therefore call it opportunistic stacking model. Concretely, the procedure duplicated all variables and inserted once a small value and once a very large value where data was initially missing. We chose biologically implausible age values of −1000 and 1000, respectively. For an illustration of the proposed model architecture, see Fig. 1 and section *Stacked-Prediction Model for Opportunistic Learning* in materials and methods for a detailed description of the model.

**Figure 1.**
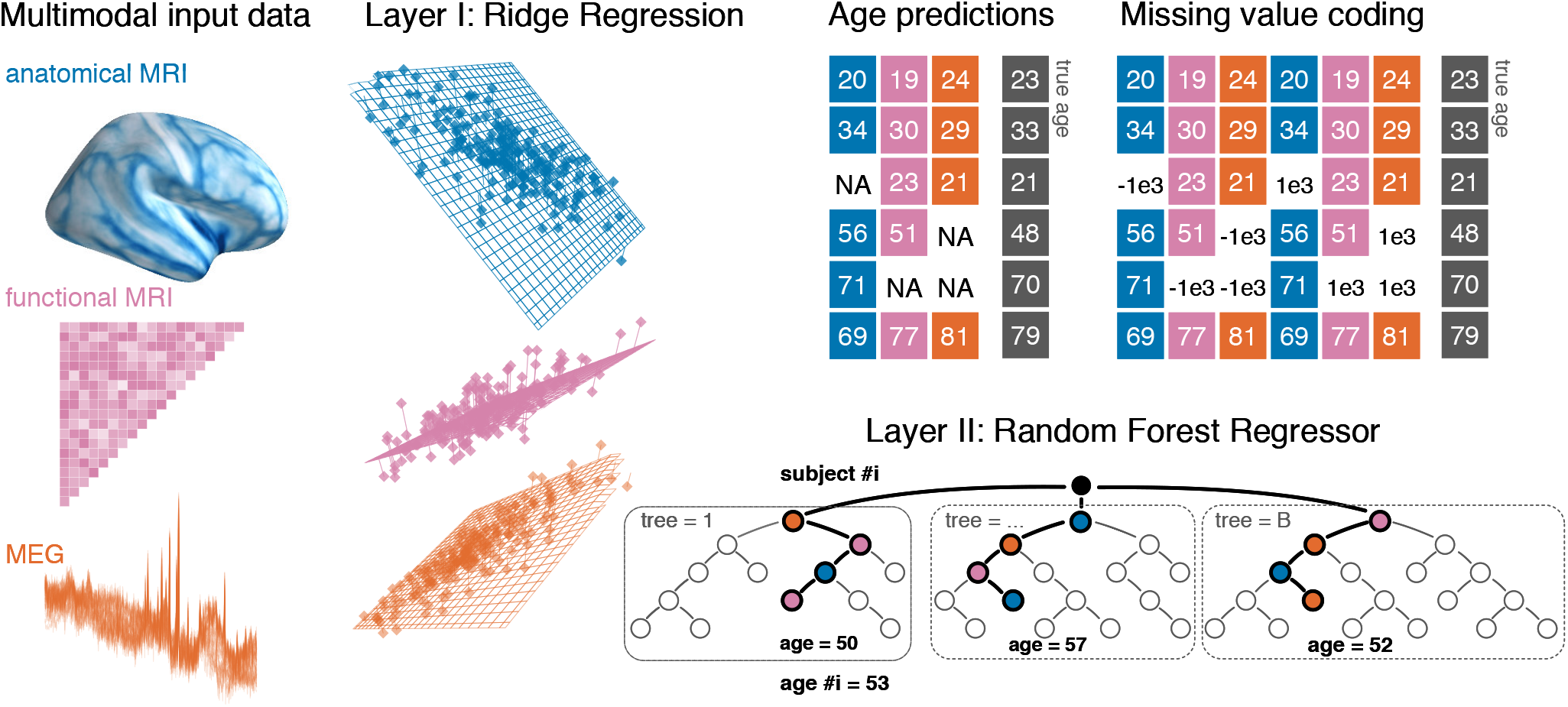
Opportunistic stacking approach. The proposed method allows to learn from any case for which at least one modality is available. The stacking model first generates, separately for each modality, linear predictions of age for held-out data. 10-fold cross-validation with 10 repeats is used. This step based on ridge regression helps reduce the dimensionality of the data by finding the major directions of variance within each modality. The predicted age is then used as derived set of features in the following steps. First, missing value are handled by a coding-scheme that duplicates the second-level data and substitutes missing values with arbitrary small and large numbers. A random forest model is then trained with the missing-value coded age-predictions from each ridge model as features to predict the actual age, potentially improving the prediction performance by combining additive information and correcting the bias of the linear model on a lower-dimensional representation.

### fMRI and MEG non-redundantly enhance anatomy-based prediction

MEG and fMRI both measure neuronal activity and convey information at smaller timescales than anatomical MRI. How would they add to the prediction of brain age when combined with anatomical MRI? Fig. 2**A** depicts a model comparison in which anatomical MRI served as baseline and which tracked changes in performance as fMRI, MEG were both added through stacking (black boxplot). Anatomical MRI scored an expected error of about 6 years, which was on average reduced by almost one year when adding either MEG or fMRI to the model. The performance gain was more than one year when adding both MEG and fMRI to the model with an expected average error about 4.7 years. The uncertainty intervals suggest that these differences were systematic and can be expected to generalize. The final drop in prediction error also suggests that MEG and fMRI carry independent information as, otherwise, the random forest would have simply picked the best of the two inputs without showing further improvement. Indeed, when comparing the prediction errors of MEG-based and fMRI-based models Fig. 2**B**, one can see that the errors are largely uncorrelated. Interestingly, fMRI, sometimes makes extreme errors for cases better predicted by MEG in younger people, whereas MEG makes errors in distinct cases from young and old age groups. When adding anatomical MRI to each model, the errors become somewhat more dependent but still showed no tight correlation.

**Figure 2.**
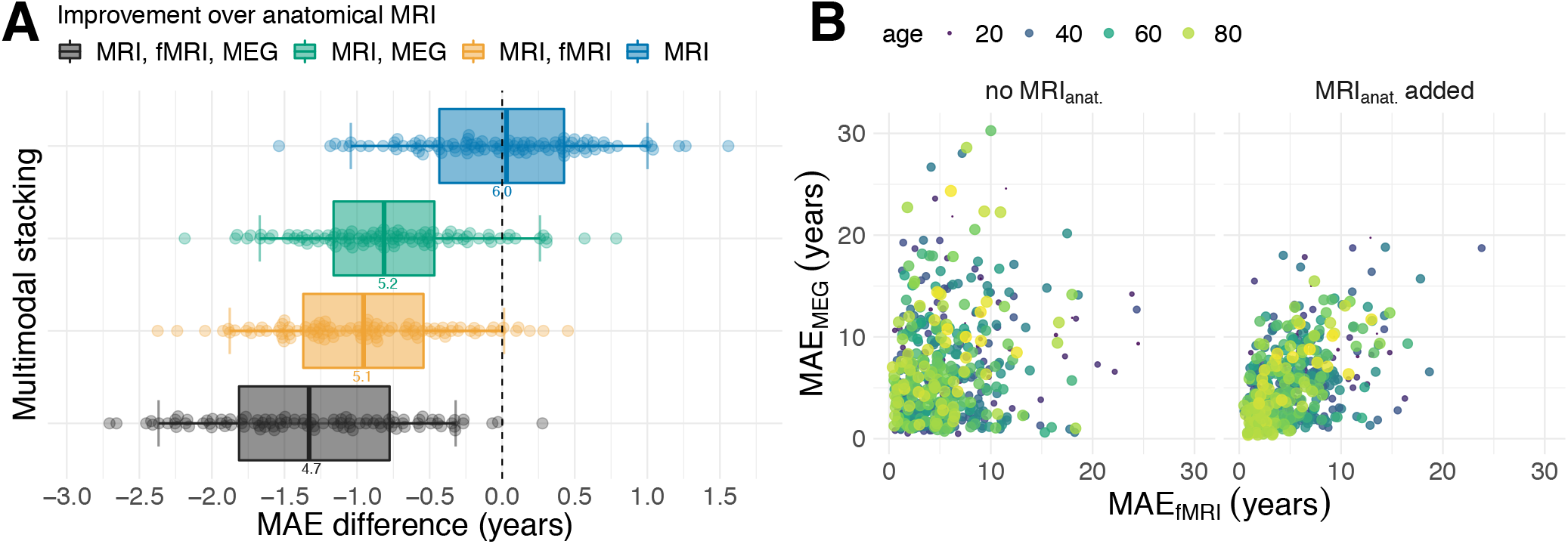
Multimodal age-prediction with MRI, fMRI and MEG. (**A**) Distribution of paired differences across cross-validation splits between stacking with anatomical MRI (blue) functional modalities, i.e., fMRI (yellow) and MEG (green) and complete stacking (black). Boxplot whiskers indicate the area including 95 percent of the values. fMRI and MEG show similar improvements over purely anatomical MRI around 0.8 years of error. Combining all modalities reduced the error by more than one year on average. (**B**) Relationship between prediction errors from fMRI and MEG. Left: unimodal models. Right: models including anatomy. Each point shows the score for one subject averaged across splits. The actual age of the subject is represented by the color and size of the dots. MEG and fMRI errors were not obviously associated. When anatomy was excluded, extreme errors occurred in different age groups. The findings suggest that fMRI and MEG convey non-redundant information. For supporting results, see *Figure 2 supplement 1-2*.

This additive component should become apparent when considering predictive simulations on how the model actually combined MEG, fMRI and MRI. *Figure 2 supplement 1* depicts a two-dimensional partial dependence analysis (Karrer et al., 2019; Hastie et al., 2005, chapter 10.13.2). Intuitively, for our model, this analysis shows how stacked predictions change as the input predictions from different modalities into the stacking layer change, two at a time. The results show that additive patterns dominate where the final age output increases equally as both input predictions increase. It is, however, noteworthy that the range of output ages was somewhat wider when the age input fMRI was manipulated, suggesting that the model trusted fMRI more than MEG.

Finally, it is worthwhile to inspect the predictions errors in a continuous fashion across age *Figure 2 supplement 2*. To better understand the impact of stacking we also included the other single-modality models (top-row). It is striking that all models show the typical brain age bias reported in the literature consisting in underfitting very old or young sub-populations (Smith et al., 2019). However, one can see how the bias is somewhat mitigated when combining multiple modalities (bottom-row). One can also see that multimodal stacking helped avoid extreme errors beyond 20 years, hence, seemed to mitigate the impact of outliers. These findings demonstrate that MEG and fMRI both add non-redundant information to an MRI-based age-prediction model. This raises the question if this additive information also implies non-redundant associations with neuropsychological assessments.

### Brain age Δ learnt from MEG and fRMI indexes distinct cognitive functions

The brain-age Δ has been interpreted as indicator of health where positive Δ has been linked to reduced fitness or health-outcomes (Cole et al., 2015, 2018). Does improved performance through stacking strengthen the effect-sizes? Do MEG and fMRI detect non-redundant associations? Fig. 3 summarizes linear correlations between the brain age *δ* and the 38 neuropsychological scores after projecting out the effect of age (see Analysis of brain-behavior correlation and Table 4 for a detailed overview). As effect sizes can be expected to be small in the curated and healthy population of the Cam-CAN dataset, we considered classical hypothesis testing for characterizing associations. Traditional significance testing (panel **A)** suggests that the best stacking models supported discoveries for between 20% (7) and 25% (9) of the scores. Dominating associations concerned fluid intelligence, depression, sleep quality (PSQI), systolic and diastolic blood pressure (cardiac features _1,2_), cognitive impairment (MMSE) and different types of memory performance (VSTM, PicturePriming, FamousFaces, EmotionalMemory). The model coefficients in panel **B** depict the strength and direction of association. One can see that stacking models not only tended to suggest more discoveries as their performance improves but also led to stronger effect sizes. However, the trend is not strict as fMRI seemed to support unique discoveries that disappeared when including the other modalities. Similarly, some effect sizes are even slightly stronger in sub-models, e.g., for fluid intelligence in MRI & MEG. A priori, the full model enjoys priority over the submodels as its expected generalization estimated with cross-validation was lower. This would imply that some of the discoveries suggested by fMRI may suffer from overfitting, but are finally corrected by the full model. Nevertheless, many of the remaining associations were found by multiple methods (e.g. fluid intelligence) whereas others were uniquely contributed by fMRI (e.g. depression) or MEG (visual short term memory) or only appear when combining all methods (sleep quality assessed by PSQI). It is also noteworthy that the directions of the effects are consistent with the predominant interpretation of the brain age Δ as indicator of mental or physical fitness (note that high PSQI score indicate sleeping difficulties whereas lower MMSE scores indicate cognitive decline) and directly confirm previous findings (Liem et al., 2017; Smith et al., 2019).

**Figure 3.**
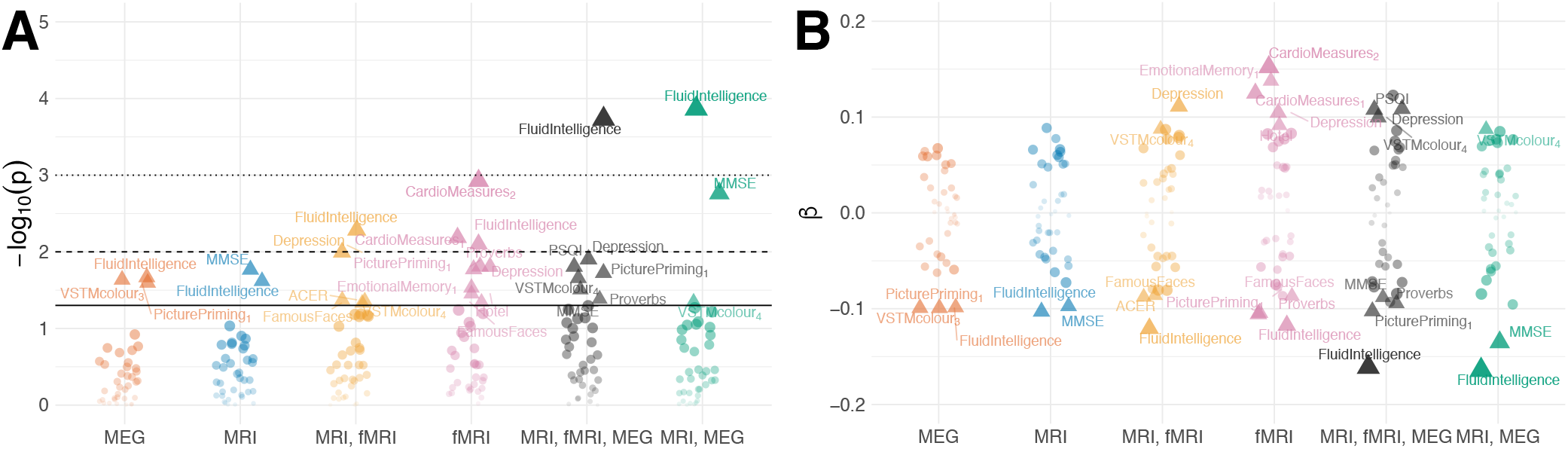
Residual correlation between brain age Δ and neuropsycholgical assessment. (**A**) Manhattan plot for linear fits of 38 neuropsychology scores against brain ageΔfrom different models (see Table 4 for overview). Y-axis: −*log*_10_(*p*). X-axis: individual scores, grouped and colored by stacking model. Arbitrary jitter is added along the x-axis to avoid overplotting. For convenience, we labelled the top scores arbitrarily thresholded by the uncorrected 5%significance level, indicated by pyramids. Fororientation, traditional 5%, 1% and 0.1% significance levels are indicated by solid, dashed and dotted lines, respectively. (**B**) Corresponding standardized coefficients ofeachlinearmodel (y-axis). Identical labelling as in **A**. One can see that, stacking often improves effect sizes for many neuropsychological scores and that different input modalities show complementary associations. For supporting results, see *Figure 3 supplement 1-3*.

These findings suggest that brain age *Δ* learnt from fMRI or MEG carries non-redundant information on clinically relevant markers of cognitive health and that combining both fMRI and MEG with anatomy can help detect health-related issues in the first place. This raises the question of what aspect of the MEG signal contributes most.

### MEG-based age-prediction is explained by source-topography of power spectra

Whether MEG or EEG-based assessment is practical in the clinical context also depends on what types of signatures are most predictive and how easily they can be combined. We therefore additionally considered purely MEG-based age prediction with the stacking method to address the following questions. Can the stacking method also be helpful to combine MEG-specific features? Are certain frequency bands of dominating importance? Is information encoded in topographic patterns or more related to interactions between brain regions? Fig. 4**A** compares alternative MEG-based models that stacked different blocks of MEG-features. All stacking models performed consistently better than chance as assessed by the error obtained when predicting the average age from the training data. Sensor space features showed the lowest performance with an expected mean absolute error around 11 years. All source space models performed markedly better with expected errors between 8 and 6.5 years. Interestingly, models based on power spectra (‘power’) only performed somewhat better than models based on bivariate connectivity (‘connectivity’), while best results were obtained when combining power with connectivity (‘combined’). Adding sensor space features did not lead to any visible improvements (‘full’). This suggests that regional changes in power spectra contain most information while connectivity adds another portion of independent information but is otherwise redundant with power. A similar picture emerges when inspecting the full model in terms of permutation-based variable importance Fig. 4 **B**. Sensor space features were least influential, whereas top contributing features were all related to power and connectivity, which, upon permutation, increased the error by up to one year. The most informative input to the stacking model were ridge regression models based on either signal power or Hilbert analytic signal power concatenated across frequency bands P_cat_, E_cat_. Additional contributions were related to power envelope connectivity (without source leakage correction) as well as source power in the beta (15-30Hz) and alpha (8-15Hz) band frequency range. The results suggest that regional cross-frequency effects are best summarized with a single linear model but additional non-linear additive effects exist in specific frequency bands.

**Figure 4.**
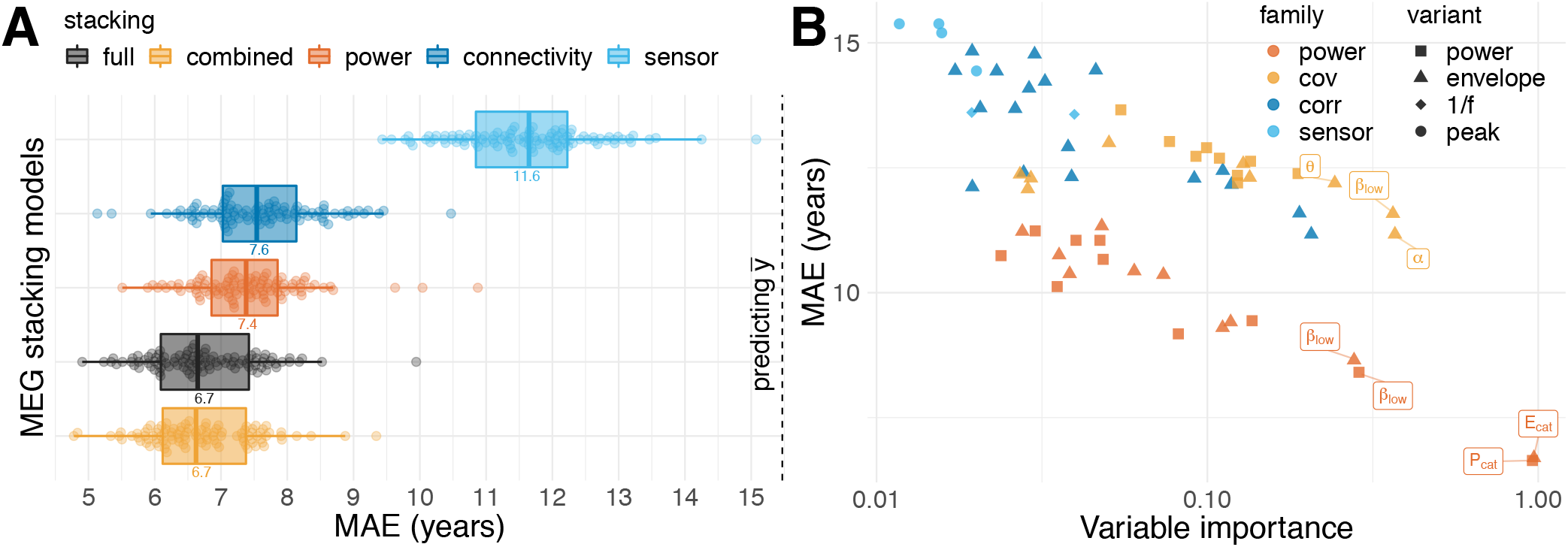
MEG-based age-prediction. (**A**) Error distributions for stacking models selected from five families of linear inputs: sensor space (sky blue), source power (dark orange), source connectivity (blue), source power and connectivity combined (light orange) and the full model based on all linear inputs. Best results were obtained when combining source power with connectivity. (**B**) Importance of linear-inputs during stacking. X-axis: mean importance (1000 permutations). Y-axis: corresponding performance of linear model. Model family is indicated by color, variants by shape. Power and covariance features were extracted for source power and envelopes. Sensor-space features encompassed 1/f and peak variants (See Table 3). Top-performing age-predictors are labeled for convenience (P = power, E = envelope, cat = concatenated across frequencies, greek letters indicate the frequency band). Some age-inputs were useful to the Random Forest despite not making strong stand-alone models. For supporting results, see *Figure 4 supplement 1-2*.

To explore how the stacking model combined the different prediction inputs, we considered a partial dependence analysis (Karrer et al., 2019; Hastie et al., 2005, chapter 10.13.2) in *Figure 4 supplement 1*. For our model, this amounts to simulating how final stacked predictions change as age predictions from the first layer linear models increase. Results revealed a staircase pattern suggesting dominant monotonic but not non-linear relationship. Moreover, the analysis revealed that more important input models had wider ranges of age predictions and were, on average, less strongly corrected by shrinkage toward the mean age. This provides some insight on how the stacking model actually improves over the linear model, that is, by pulling implausible extreme predictions towards the mean prediction. Importantly, the best stacked models scored lower errors than the best linear models (*Figure 4 supplement 2*), suggesting that stacking achieved more than mere variable selection and instead extracted non-redundant information from the inputs.

These findings show that MEG-based prediction of age, is enabled by features that can be relatively easily accessed in terms of computation and data processing. Moreover, the stacking approach applied to MEG data helped to improve beyond the linear model.

### Advantages of multimodal stacking can be maintained on populations with incomplete data

One important obstacle for combining signals from multiple modalities in clinical settings is that not all modalities are available for all cases. So far we have restricted the analysis to 536 cases for which all modalities were present. Can the advantage of multimodal stacking be maintained in the absence of complete data or will missing values mitigate prediction performance? To investigate this question, we trained our stacked model on all 674 cases for which we had the opportunity to extract at least one feature on any modality, hence, we termed it opportunistic stacking (see 1 and Table 1 in section Sample in materials and methods). We first compared the opportunistic model with the restricted model on the cases with complete data Fig. 5**A**. Across stacking models, performance was virtually identical, even when extending the comparison to the cases available to the sub-model with fewer modalities, e.g., MRI & fMRI. We then scored the fully opportunistic model trained on all cases and all modalities and compared it to different non-opportunistic sub-models on restricted cases (Fig. 5**A**, squares). The fully opportunistic model always out-performed the sub-model. This raises the question of how the remaining cases would be predicted for which fewer modalities were available. Fig. 5**B** shows the performance of the opportunistic split by sub-groups defined by different combinations of input modalities available. As expected, performance degraded considerably on sub-groups for which important features (as delineated by the previous results) were not available. See for example the sub-group for which only sensor-space MEG was available. This is not surprising, as prediction has to be based on data and is necessarily compromised if features important at train-time are not available at predict-time. Importantly however, this finding suggests that the opportunistic model operates conservatively: The performance on the sub-groups reflects the quality of the features available, hence, enables learning from the entire data.

**Figure 5.**
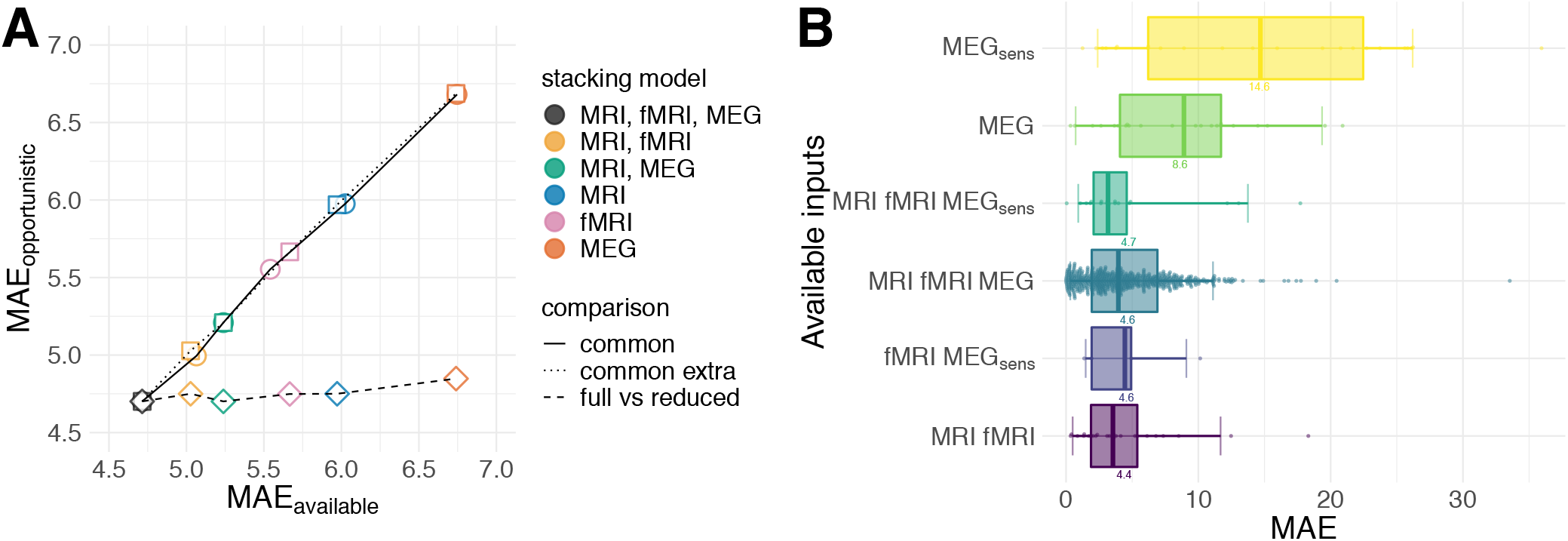
Opportunistic learning performance. (**A**) Comparisons between opportunistically trained model and models restricted to common available cases. Opportunistic versus restricted model with different combinations scored on all 536 *common* cases (circles). Same analysis extended to include *extra common* cases available for sub-models (squares). Fully opportunistic stacking model (all cases, all modalities) versus reduced non-opportunistic sub-models (fewer modalities) on the cases available to the given sub-model (diamonds). One can see that multimodal stacking is generally of advantage whenever multiple modalities are available and does not impact performance compared to restricted analysis on modality-complete data. (**B**) Performance for opportunistically trained model for subgroups defined by different combinations of available input modalities, ordered by average error. Points depict single-case prediction errors. Boxplot-whiskers show the 5% and 95% uncertainty intervals. When performance was degraded, important modalities were absent or the number of cases was small.

**Table 1.** Available cases by input modality

| modality | MEG sensor | MEG source | MRI | fMRI | common cases |
| --- | --- | --- | --- | --- | --- |
| cases | 589 | 600 | 621 | 626 | 536 |

*Note.* MEG sensor space cases reflect separate task-related recordings. MEG source space cases are based on the resting state recordings.

## Discussion

We have demonstrated improved learning of surrogate biomarkers by combining electrophysiology, functional and anatomical MRI. Here, we have focused on the example of age-prediction by multimodal modeling on 674 subjects from the Cam-CAN dataset, the currently largest publicly available collection of MEG, fMRI and MRI data. Our results suggest that MEG and fMRI both substantially improved age-prediction when combined with anatomical MRI. We have then explored potential implications of the ensuing brain-age Δas a surrogate-biomarker for cognitive and physical health. Our results suggest that MEG and fMRI convey non-redundant information on cognitive functioning and health, e.g., fluid intelligence, memory, sleep quality, cognitive decline and depression. Moreover, combining all modalities has led to lower prediction errors. Inspection of the MEG-based models suggested unique information on aging is conveyed by regional distribution of power in the *α* (8-12Hz) and *β* (15-30Hz) frequency bands, in line with the notion of spectral finger prints (Keitel and Gross, 2016). When applied in clinical settings, multimodal approaches should make it more likely to detect brain-behavior associations. We have, therefore, addressed the issue of missing values, which is an important obstacle for multimodal learning approaches in clinical settings. Our stacking model, trained on the entire data with an opportunistic strategy, performed equivalently to the restricted model on common subsets of the data and helped exploiting multimodal information to the extent available. This suggests, that the advantages of multimodal prediction can be maintained in practice.

### fMRI and MEG reveal complementary information on cognitive aging

Our results have revealed complementary effects of anatomy and neurophysiology in age-prediction. When adding either MEG or fMRI to the anatomy-based stacking model, the prediction error markedly dropped (Fig. 2**A**). Both, MEG and fMRI helped gain almost one year of error compared to purely anatomy-based prediction. This finding suggests that both modalities access equivalent information. This is in line with a recent literature on correspondence of MEG with fMRI in resting state networks, highlighting the importance of spatially correlated slow fluctuations in brain oscillations (Hipp and Siegel, 2015; Hipp et al., 2012; Brookes et al., 2011) and, more specifically, a recent finding suggesting that age-related variability in fMRI and EEG is independent to a substantial degree (Kumral et al., 2019).

Interestingly, the prediction errors of models with MEG and models with fMRI were not systematically correlated (Fig. 2**B**, left panel). In some subpopulations, they even seemed anti-correlated, such that predictions from MEG or fMRI, for the same cases, were either accurate or extremely inaccurate. This additional finding would actually suggest that the improvements of MEG and fMRI over anatomical MRI are not due to shared information but due to their access to complementary information that helps predicting distinct cases. Indeed, when we combined MEG and fMRI in one common stacking model together with anatomy, performance, improved on average by 1.3 years over the purely anatomical model, which is almost half a year more precise than the previous MEG-based and fMRI-based models.

The results strongly suggest the presence of an additive component, in line with the common intuition that MEG and fMRI are complementary with regard to spatial and temporal resolution. In this context, our results on performance decomposition in MEG (Fig.4) delivers one potentially interesting hint. The source topography of power spectral density, especially in the *α*(8 − 15*Hz*) and *β*(15 − 26*Hz*) range turned out to be the single most contributing type of feature (Fig. 4**A**).

However, connectivity features, in general, and power-envelope connectivity, in particular, contributed substantively but rather weakly (Fig; 4**B**, *Figure 4 supplement*). Interestingly, applying orthogonalization (Hipp et al., 2012; Hipp and Siegel, 2015) for removing source leakage did not visibly improve performance (*Figure 4 supplement 2*). Against the background of MEG-fMRI correspondence, which has highlighted the importance of slow fluctuations of brain rhythms (Hipp and Siegel, 2015; Brookes et al., 2011), this finding suggests that what renders MEG non-redundant with regard to fMRI are regional differences in the balance of fast brain-rhythms, in particular in the *α* − *β* range. If this turned out to be true, one could expect that electrophysiology will make a true additive contribution to prediction problems in which fast brain rhythms are strongly statistically related to the target.

### Brain age Δ as sensitive index of normative aging

In this study we have conducted an exploratory analysis on what might be the cognitive and health-related implications of our prediction models. Our findings suggest the brain age Δ shows substantive associations with about 20-25% of all neuropsychological measures included. The overall big-picture is congruent with the brain age literature (see discussion in Smith et al. 2019 for an overview) and supports the interpretation of the brain age Δ as index of decline of physical health, well-being and cognitive fitness. In this sample, larger values of the Δ were globally associated with elevated depression scores, higher blood pressure, lower sleep quality, lower fluid intelligence, lower scores in neurological assessment and lower memory performance. Most strikingly, we found that fMRI and MEG contribute unique discoveries, even when combined (Fig. 3). For example, the association with depression appeared first when predicting age from fMRI. On the other hand, visual short term memory appears first in MEG-based models. Moreover, the association with fluid intelligence only manifested itself when including MEG. Finally, sleep quality emerged once all modalities were combined. This extends the previous discussion in suggesting that MEG and fMRI are not only complementary for prediction but also with regard to characterizing brain-behavior mappings. Moreover, it is enticing to speculate that the regional power of fast-paced *α* and *β* band brain rhythms allows one to capture fast-paced components of cognitive processes such as attentional sampling or adaptive attention (Gola et al., 2013; Clark et al., 2004), which, in turn might explain unique variance in certain cognitive facets, such as fluid intelligence (Ouyang et al., 2019) or visual short-term memory (Tallon-Baudry et al., 2001). On the other hand, functional connectivity between cortical areas and subcortical structures, in particular the hippocampus, may be key for depression and is well captured with fMRI (Stockmeier et al., 2004; Sheline et al., 2009; Rocca et al., 2015). Unfortunately, modeling such mediation effects exceeds the scope of the current work, although it would be worth being tested in an independent study with a dedicated design.

However, it is important to appreciate these findings carefully. One could argue that the overall effect sizes were too low to be considered practically interesting. Indeed, the strength of linear association was below 0.5 in units of standard deviations of the normalized predictors and the target. On the other hand, it is important to consider that the Cam-CAN sample consists of healthy individuals only. It appears, thus, as rather striking that systematic and neuropsychologically plausible effects can be detected. The finding, therefore, argues in favor of the brain age Δ being a sensitive marker of normative aging. The effects are expected to be far more pronounced when applying the method in clinical settings, i.e., in patients suffering from mild cognitive impairment, depression, neurodevelopmental or neurodegenerative disorders. This suggests that brain age Δ might be used as a screening tool for a wide array of clinical settings for which the Cam-CAN dataset could serve as a normative sample.

### Translation to the clinical setting

One critical factor for application of our approach in the clinic is the problem of incomplete availability of medical imaging and physiological measurements. Here, we addressed this issue by applying an opportunistic learning approach which enables learning from the data available at hand. Our analysis of opportunistic learning applied to age prediction revealed viable practical alternatives to confining the analysis to cases for which all measurements are available. In fact, adding extra cases with incomplete measurements never harmed prediction of the cases with complete data and the full multimodal stacking always outperformed sub-models with fewer modalities (Fig. 5**A**). Moreover, the approach allowed maintaining and extending the performance to new cases with incomplete modalities (Fig. 5**B**).Importantly, performance on such subsets was explained by the performance of a reduced model with the remaining modalities. Put differently, opportunistic stacking performed as good as a model restricted to data with all modalities. Practically speaking, the approach allows one to improve predictions case-wise by including electrophysiology next to MRI or MRI next to electrophysiology, whenever there is the opportunity to do so.

A second critical factor for translating our findings into the clinic is that most of the time, it is not high-density MEG that is available but low-density EEG. In this context, our finding that the source-topography power spectrum was the mostimportantfeature is of clear practical interest. This is because it suggests that a rather simple statistical object accounts for the bulk of the performance of MEG. The topography of power spectra can be computed on any multichannel EEG device in a few steps and only yields, per frequency band, as many variables as there are channels. Moreover, from a statistical standpoint, computing the power spectrum amounts to estimating the marginal expectation of the signal variance, which can be thought of as main effect. On the other hand, connectivity is often operationalized as bivariate interaction, which gives rise to a more complex statistical object of higher dimensionality whose precise, reproducible estimation may require far more samples. Moreover, as is the case for power envelope connectivity estimation, additional processing steps each of which may add researcher degrees of freedom (Simmons et al., 2011), such as the choice between Hilbert (Brookes et al., 2011) versus Wavelet filtering (Hipp et al., 2012), types of orthogonalization (Baker et al., 2014), and potentially thresholding for topological analysis (Khan et al., 2018). This nourishes the hope that our findings will generalize and similar performance can be unlocked on simpler EEG devices with fewer channels. While clinical EEG may not well resolve functional connectivity it may be good enough to resolve changes in the source geometry of the power spectrum. On the other hand, source localization may be critical in this context as linear field spread actually results in a non-linear transform when considering the power of a source (Sabbagh et al., 2019a,b). Indeed, our model has strongly favored source-level features. However, in practice, it may be hard to conduct high-fidelity source localization on the basis of low-density EEG and frequently absent information on the individual anatomy. It will therefore be critical to benchmark and improve learning from power topographies in clinical settings (Sabbagh et al., 2019a).

Finally, it is worthwhile to highlight that here we have focused on age, in the more specific context of the brain age Δ surrogate biomarker in order to be able to benefit from a relatively large benchmark dataset. However, the proposed approach is fully compatible with any target of interest and may be easily applied directly to clinical end points, e.g., drug dosage, survival or diagnosis. Moreover, the approach presented here can be easily adapted to work with classification problems, for instance, by exchanging ridge regression with logistic regression and using a random forest classifier in the stacking layer. We have provided all materials from our study in form of publicly available version-controlled code with the hope to help other teams of biomedical researchers to adapt our method to their prediction problem.

## Materials and Methods

### Sample

Here, we included MEG(task & rest), fMRI (rest), MRI and neuropsychological data(cognitive tests, home-interview, questionnaires) from the CAM-Can dataset (Shafto et al., 2014). Our sample comprised 674 (340 female) healthy individuals between 18 (female = 18) to 88 (female = 87) years with an average of 54.2 (female = 53.7) and a standard deviation of 18.7 (female = 18.8) years. We included data according to availability and did not apply an explicit criterion for exclusion. When automated processing resulted in errors, we did not manually repair the computation. This induced additional missing data for some cases. A summary of available cases by input modality is reported in Table 1 in the appendix. For technical details regarding the MEG, fMRI, and MRI data acquisition, please consider the Cam-CAN reference publications (Shafto et al., 2014; Taylor et al., 2017).

### Feature extraction

For MEG, we analyzed sensor space features related to timing, peak frequency and temporal autocorrelation, and source space features related to the regional power in nine frequency bands, power envelopes and bivariate interactions. The definition of frequency bands (see Table 2) was adopted from the Human Connectome Project (Larson-Prior et al., 2013). In general, the selection of features was guided by the literature on aging-related EEG and MEG signatures. More specifically, we wanted to enable more targeted comparisons between MEG and fMRI by including power envelopes, i.e., the slow fluctuations of power and their bivariate correlations between them. These have been shown repeatedly to give rise to spatial patterns that correspond to fMRI resting state networks. On the other hand, we wanted to exploit the potentially unique capacity of the MEG to access topographic information induced by fast-paced brain rhythms emerging from regional sources. We therefore included source power and covariance among the features. To mitigate distortions of the non-linear source power through the individual anatomy (forward model) we used source localization. For MRI and fMRI, we adapted the approach established by Liem et al. (2017) and focused on cortical thickness, cortical surface area and subcortical volumes. For fMRI, we computed bivariate functional connectivity estimates. An overview on all features used is presented in Table 3. In the following, we describe computation details.

**Table 2.** Frequency band definitions

| name | low | $\delta$ | $\theta$ | $\alpha$ | $\beta_1$ | $\beta_2$ | $\gamma_1$ | $\gamma_2$ | $\gamma_3$ |
| --- | --- | --- | --- | --- | --- | --- | --- | --- | --- |
| range (Hz) | 0.1 – 1.5 | 1.5 – 4 | 4 – 8 | 8 – 15 | 15 – 26 | 26 – 35 | 35 – 50 | 50 – 74 | 76 – 120 |

### MEG features

#### peak evoked latency

Sensory processing may slow down in the course of aging (Price et al., 2017). Here, we assessed the evoked response latency during auditory, visual and simultaneous audiovisual stimulation (index 1, Table 3). For each of the conditions, we first computed the evoked response. Then, we computed the root-mean-square across gradiometers and looked up the time of the maximum. This yielded in total three latency values.

**Table 3.** Summary of extracted features.

| # | Modality | Feature | Reduction | Variants | Family |
| --- | --- | --- | --- | --- | --- |
| 1 | MEG | peak evoked latency | max | aud, vis, audvis | sensor |
| 2 | ... | $\alpha$ peak | max | | ... |
| 3 | ... | 1/f slope | channels | low, $\gamma$ | ... |
| 4 | ... | power | 448 ROIs | low, $\delta, \theta, \alpha, \beta_{1,2}, \gamma_{1,2,3}$ | power |
| 5 | ... | power envelope | ... | ... | ... |
| 6 | ... | covariance (cov.) | ... | ... | connectivity |
| 7 | ... | envelope (env.) cov. | ... | ... | ... |
| 8 | ... | env. correlation (corr.) | ... | ... | ... |
| 9 | ... | env. corr. adjusted | ... | ... | ... |
| 10 | fMRI | correlation | 256 ROIs |  | ... |
| 11 | MRI | cortical (cort.) thickness | 5124 vertices |  | anatomy |
| 12 | ... | cort. surface area | 5124 vertices |  | ... |
| 13 | ... | subcortical volumes | 66 ROIs |  | ... |

#### α-band peak frequency

Research suggests that the alpha-band frequency may be lower in older people. Here, we computed the resting-state power spectrum using a Welch estimator (index 2, Table 3). Then, we estimated the peak frequency between 6 and 15 Hz on occipito-parietal magnetometers after removing the 1/f trend using a polynomial regression (degree = 15) by computing the maximum power across sensors and looking up the frequency bin. This yielded one peak value per subject.

#### 1/f slope

Long-range auto-correlation in neural time-series gives rise to the characteristic 1/f decay of power on a logarithmic scale. Increases of neural noise during aging are thought to lead to reduced autocorrelation, hence a more shallow slope (Voytek et al., 2015). We computed the 1/f slope from the Welch power spectral estimates above on all magnetometers using linear regression (index 3, Table 3). The slope is given by the 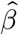 of the linear fit with the logarithm of the frequencies as predictor and the log power as target. We obtained one estimate for each of the 102 magnetometers, resulting in a 1/f topography. No further reduction was applied.

#### source power and connectivity

The cortical generators of the brain-rhythms dominating the power spectrum change across life-span. To mitigate geometrical distortions through individual anatomy, we used source-localization to estimate the topography of power. We used a subdivision of the Desikan-Killiany atlas (Desikan et al., 2006) that comprised 448 ROIs (Khan et al., 2018). We bandpass-filtered signals into frequency bands (see Table2), computed minimum norm source-estimates and then summarized the source-time courses ROI-wise by the first principal components. We then computed the covariance matrix and used as power estimates the 448 diagonal entries (index 4 Table3). The off-diagonal entries served as connectivity estimates. Covariance matrices live in a non-Euclidean curved space. To avoid model violations at subsequent modeling stages, we used tangent space projection (Varoquaux et al., 2010) to vectorize the lower triangle of the covariance matrix. This projection allows one to treat entries of the correlation matrix as regular Euclidean objects. This yielded 448 × 448/2-(448/2) = 100, 128 connectivity values(index 6 Table3).

#### source power envelopes and connectivity

Brain-rhythms are not constant in time but fluctuate in intensity. These slow fluctuations are technically referred to as power envelopes and may show characteristic patterns of spatial correlation. To estimate power envelopes, for each frequency band, we computed the analytic signal using the Hilbert transform. We applied the same procedure as for source power (paragraph above) to estimate the source power of the envelopes (index 5, Table 3) and their connectivity. In the MEG literature, envelope correlation is a well established research topic. We therefore also computed, beyond the covariance, the commonly used normalized Pearson correlations and orthogonalized Pearson correlations which are designed to mitigate source leakage (index 7-9, Table 3). However, as a result of orthogonalization, the resulting matrix is not any longer positive definite, hence, cannot be projected to the tangent space. We therefore used Fisher’s Z-transform (Silver and Dunlap, 1987) to convert the correlation matrix into a set of standard-normal variables. The transform is defined as the inverse hyperbolic tangent function of the correlation coefficient: 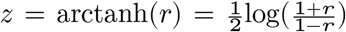. This yielded 448 power envelope power estimates and 100,128 connectivity values per estimator.

### fMRI features

#### functional connectivity

Large-scale neuronal interactions between distinct brain networks has been repeatedly shown to change during healthy aging. To estimate macroscopic functional connectivity, we used the MODL atlas with 256 functional ROI (Mensch et al., 2016). We then computed bivariate amplitude interactions using Pearson correlations from the ROI-wise average time-series (index 10, Table. 3). Again, we used tangent space projection (Varoquaux et al., 2010) to vectorize the correlation matrices. This yielded 32,640 connectivity values from the lower triangle of each matrix. No further reduction was applied.

### MRI features

#### cortical thickness

Aging-related brain atrophy has been related to thinning of the cortical tissue, e.g., (Thambisetty et al., 2010). Here, we extracted cortical thickness estimates on from the Freesurfer (Fischl, 2012) segmentation on a grid of 5,124 vertices in `fsaverage4` space obtained from the `mris_preproc` script (index 11, Table 3). No further reduction was computed.

#### cortical surface area

Aging is also reflected in shrinkage of the cortical surface itself, e.g., (Lemaitre et al., 2012). Hence, we also extracted cortical surface area estimates on from the Freesurfer segmentation on a grid of 5,124 vertices in `fsaverage4` space obtained from the `mris_preproc` script (index 12, Table 3). No further reduction was computed.

#### subcortical volumes

The volume of subcortical structures has been linked to aging (Murphy et al., 1992). Here, we used the `asegstats2table` to obtain estimates of the subcortical volumes and global volume, yielding 66 values for each subject with no further reductions (index 13, Table 3).

### Stacked-Prediction Model for Opportunistic Learning

We used the stacked-prediction framework (Wolpert, 1992) to build our predictive model. However, we made the important specification that input models were regularized linear models trained on different blocks of variables and block-wise stacking of predictions was achieved by a local, non-linear regression model. Our model can be intuitively denoted as follows:

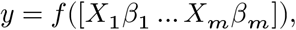

Here, each *X_j_β_j_* is the vector of predictions *ŷ_j_* of the target vector *y* from the j*th* model fitted using input data *X_j_*:

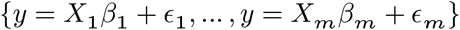

Here, we used ridge regression as input model and a random forest regressor as general function approximator *f* [Chp. 15.4.3](Hastie et al., 2005). A visual illustration of the model is presented in Fig. 1.

#### Input Layer: Ridge Regression

Results from statistical decision theory suggests that, for linear models, the expected out-of-sample error increases only linearly with the number of variables included in a prediction problem (Hastie et al., 2005, chapter 2), not exponentially. In practice, biased (or penalized) linear models with Gaussian priors on the coefficients, i.e., ridge regression (or logistic regression for classification) with *ℓ*_2_-penalty (squared *ℓ*_2_ norm) are hard to outperform in neuroimaging settings (Dadi et al., 2019). Ridge regression can be seen as extension of ordinary least squares (OLS) where the solution is biased such that the coefficients estimated from the data are conservatively pushed towards zero:

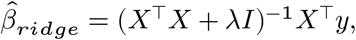

The estimated coefficients approach zero as the penalty term λ grows, and the solution approaches the OLS fit as λ gets closer to zero. This is the same as assuming that the coefficient vector comes from a Gaussian distribution centered around zero [chapter 7.3](Efron and Hastie, 2016):

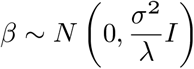

In practice, reasonable priors are often unknown, hence, λ is chosen in a data-driven fashion such that one improves the expected out-of-sample error, e.g., tuned using cross-validation. We tuned the λ using generalized cross-validation (Golub et al., 1979) and considered 100 candidate values on a logarithmic scale between 10^-3^ and 10^5^.

#### Stacking Layer: Random Forest Regression

However, the performance of the ridge model in high dimensions comes at the price of increased bias. The stacking model tries to alleviate this issue by reducing the dimensionality in creating a derived data set of linear predictions, which can then be forwarded to a more flexible local regression model. Here, we chose the random forest algorithm (Breiman, 2001) which can be seen as a general function approximator and has been interpreted as adaptive nearest neighbors algorithm (Hastie et al., 2005, chapter 15.4.3). Random forests can learn a wide range of functions and are capable of automatically detecting non-linear interaction effects with little tuning of hyper-parameters. They are based on two principles: regression trees and bagging (bootstrapping and aggregating). Regression trees are non-parametric methods and recursively subdivide the input data by finding combinations of thresholds that relate value ranges of the input variables to the target. The principle is illustrated at the right bottom of Fig. 1. For a fully grown tree, each sample falls into one leaf of the tree which is defined by its unique path through combinations of input-variable thresholds through the tree. However, regression trees tend to easily overfit. This is counteracted by randomly generating alternative trees from bootstrap replica of the dataset and randomly selecting subset of variables for each tree. Importantly, thresholds are by default optimized with regard to a so-called impurity criterion such as entropy or mutual information. Predictions are then averaged, which mitigates overfitting and also explains how thresholds can lead to continuous predictions.

In practice, it is common to use a generous number of trees as performance plateaus once a certain number is reached, which may lay between hundreds or thousands. Here, we used 1000 trees. Moreover, limiting the overall depth of the trees can increase bias and mitigate overfitting at the expense of model complexity. An intuitive way of conceptualizing this step is to think of the tree-depth in terms of orders interaction effects. A tree with three nodes enables learning three-way interactions. Here, we tuned the model to choose between depth-values of 4, 6, or 8 or the option of not constraining the depth. Finally, the total number of features sampled at each node determines the degree to which the individual trees are independent or correlated. Small number of variables decorrelate the trees but make it harder to find important variables as the number of input variables increases. On the other hand, using more variables at once leads to more exhaustive search of good thresholds, but may increase overfitting. As our stacking models had to deal with different number of input variables, we had to tune this parameter and let the model select between 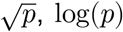 and all *p* input variables. We implemented selection of tuning-parameters by grid search as (nested) 5-fold crossvalidation. For performance quantification, we used the mean absolute error.

#### Stacked cross-validation

We used a 10-fold cross-validation scheme. To mitigate bias due to the actual order of the data, we repeated the procedure 10 times while reshuffling the data at each repeat. We then generated age-predictions from each Input-layer model on the left-out folds, such that we had for each case one age-prediction per repeat. We then stored the indices for each fold to make sure the random forest was trained on left-out predictions for the ridge models. This ensured that the input-layer train-test splits where carried forward to the stacking-layer and that the stacking model was always evaluated on left-out folds in which the input ages are actual predictions and the targets have not been seen by the model.

Here, we customized the stacking procedure to be able to unbox and analyze the input-age predictions and implement opportunistic handling of missing values.

#### Variable importance

Regression trees are often inspected by estimating the impact of each variable on the prediction. This is commonly achieved by computing the so-called variable importance. The idea is to track and sum across all trees the reduction of impurity each time a given variable is used to split. However, it has been shown that in correlated trees, variable importance can be biased and lead to masking effects, i.e., fail to detect important variables (Louppe et al., 2013) or suggest noise-variables to be important. One potential remedy is to increase the randomness of the trees, e.g., by selecting randomly a single variable for splitting and using extremely randomized trees (Geurts et al., 2006; Engemann et al., 2018), as it can be mathematically guaranteed that in fully randomized trees only actually important variables are assigned importance (Louppe et al., 2013). However, such measures may mitigate performance. Here, we used an alternative, model-agnostic approach, which consists in permuting randomly one variable at a time and measuring the drop in performance at the scale of the scoring. This is approach is closely related to the method described in the original random forest paper (Breiman, 2001), with the difference that we used cross-validation instead of out-of-bag estimates. This procedure has the known disadvantage, that it does not take into account the conditional nature of variable importance. For example, a variable may not be so important in itself but its interaction with other variables makes it an important predictor. On the other hand, the permutation importance approach has the advantage that importance is intuitively expressed in units of the error scoring and that it avoids masking.

#### Opportunistic Learning with Missing Values

An important option for our stacking model concerns handling missing values. Here, we implemented the double-coding approach (Josse et al., 2019) which duplicates the features and once assigns the missing value a very small and once a very large number (see also illustration Fig. 1). As our stacked input data consisted of age predictions from the ridge models, we used biologically but also statistically implausible values of −1000 and 1000, respectively. This amounts to turning missing values into features and let the stacking-model potentially learn from the missing values, as the reason for the missing value may contain information on the target. For example, an elderly patient may not be in the best conditions for an MRI scan, but nevertheless qualifies for electrophysiological assessment.

To implement opportunistic stacking, we considered the full dataset with missing values and then kept track of missing data while training the input-layer. This yielded the stacking-data consisting of the age-predictions and missing values. Stacking was then performed after applying the feature-coding of missing values. This procedure made sure that all training and test splits were defined with regard to the full cases and, hence, the stacking model could be applied to all cases after feature-coding of missing values.

### Analysis of brain-behavior correlation

To explore the cognitive implications of the brain age Δ, we computed correlations with the neurobehavioral score from the Cam-CAN dataset. Table 4 lists the scores we considered. The measures fall into three broad classes: neuropsychology, physiology and questionnaires (‘Type’ columns in Table 4). Extraction of neuropsychological scores sometimes required additional computation, which followed the description in Shafto et al. 2014, (see also ‘Variables’ column in Table 4). For some neuropsychological tasks, the Cam-CAN dataset provided multiple scores and sometimes the final score of interest as described in Shafto et al. 2014, had yet to be computed. At times, this amounted to computing ratios, averages or differences between different scores. In other scores, it was not obvious how to aggregate multiple interrelated sub-scores, hence, we computed summaries by extracting the first principal component. In total, we included 38 variables. All neuropsychology and physiology scores (up to #17) were the scores available in the ‘cc770-scored’ folder from release 001 of the Cam-CAN dataset. We selected the additional questionnaire scores (#18-23) on theoretical grounds to provide an assessment of clinically relevant individual differences in cognitive functioning. The brain age Δ was defined as the difference between predicted and actual age of the person

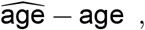

**Table 4.** Summary of neurobehavioral scores

| # | Name | Type | Variables (=38) |
| --- | --- | --- | --- |
| 1 | Benton faces | neuropsychology | total score (1) |
| 2 | Emotional expression recognition | ... | PC1 of RT (1) |
| 3 | Emotional memory | ... | PC1 by memory type (3) |
| 4 | Emotion regulation | ... | positive & negative reactivity, regulation (3) |
| 5 | Famous faces | ... | mean familiar details ratio (1) |
| 6 | Fluid intelligence | ... | total score (1) |
| 7 | Force matching | ... | Finger- & slider-overcompensation (2) |
| 7 | Hotel task | ... | time(1) |
| 9 | Motor learning | ... | M & SD of trajectory error (2) |
| 10 | Picture priming | ... | baseline RT, baseline ACC (4) |
| ... | ... | ... | M prime RT contrast, M target RT contrast |
| 11 | Proverb comprehension | ... | score (1) |
| 12 | RT choice | ... | M RT (1) |
| 13 | RT simple | ... | M RT (1) |
| 14 | Sentence comprehension | ... | unacceptable error, M RT (2) |
| 15 | Tip-of-the-tounge task | ... | ratio (1) |
| 16 | Visual short term memory | ... | K (M,precision,doubt,MSE) (4) |
| 17 | Cardio markers | physiology | pulse, systolic & diastolic pressure (3) |
| 18 | PSQI | questionnaire | total score (1) |
| 19 | Hours slept | ... | total score (1) |
| 20 | HADS (Depression) | ... | total score (1) |
| 21 | HADS (Anxiety) | ... | total score (1) |
| 22 | ACE-R | ... | total score (1) |
| 23 | MMSE | ... | total score (1) |

Note. M = mean, SD = standard deviation, RT = reaction time, PC = principal component, ACC = accuracy,
PSQI = Pittsburgh Sleep Quality Index HADS = Hospital Anxiety and Depression Scale,
ACE-R = Addenbrookes Cognitive Examination Revised, MMSE = Mini-Mental State Examination.
Numbers in parentheses indicate how many variables were extracted.

such that positive values quantify overestimation and negative value underestimation. A common problem in establishing brain-behavior correlations for brain age is spurious correlations due to shared age-related variance in the brain age Δ and the neurobehavioral score (Smith et al., 2019). To mitigate confounding effects of age, we computed the age residuals as

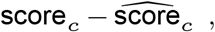

where *score_c_* is the current score and the predicted score 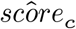 is obtained from the following polynomial regression:

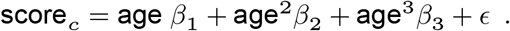

To obtain comparable coefficients across scores, we standardized both the age and the scores.

## MEG data processing

### Data Acquisition

MEG recorded at a single site using a 306 VectorView system (Elekta Neuromag, Helsinki). This system is equipped with 102 magnetometers and 204 orthogonal planar gradiometers is placed light magnetically shielded room. During acquisition, an online filter was applied between around 0.03 Hz and 1000Hz. To support offline artifact correction, vertical and horizontal electrooculogram (VEOG, HEOG) as well as electrocardiogram (ECG) signal was concomitantly recorded. Four Head-Position Indicator (HPI) coils were used to track head motion. All types of recordings, i.e., resting-state, passive stimulation and the active task lasted about 8 minutes. For additional details on MEG acquisition, please consider the reference publications of the CAM-Can (Taylor et al., 2017; Shafto et al., 2014). The following sections will describe the custom data processing conducted in our study.

### Artifact Removal

#### Environmental artifacts

To mitigate contamination of the MEG signal with artifacts produced by environmental magnetic sources, we applied temporal signal-space-separation (tSSS) (Taulu and Kajola, 2005). The method uses spherical harmonic decomposition to separate spatial patterns produced by sources inside the head from patterns produced by external sources. We used the default settings with eight components the harmonic decomposition of the internal sources, and three for the external sources on a ten seconds sliding window. We used a correlation threshold of 98% to ignore segments in which inner and outer signal components are poorly distinguishable. We performed no movement compensation, since there were no continuous head monitoring data available at the time of our study. The origin of internal and external multipolar moment space was estimated based on the head-digitization. We computed tSSS using the MNE `maxwell_filter` function (Gramfort et al., 2013) but relied on the SSS processing logfiles from Cam-CAN for defining bad channels.

#### Physiological artifacts

To mitigate signal distortions caused by eye-movements and heart-beats we used signal space projection (SSP) (Uusitalo and Ilmoniemi, 1997). This method learns principal components on contaminated data-segments and then projects the signal into the sub-space that is not correlated with the artifact. To obtain clean estimates, we excluded bad data segments from the EOG/ECG channels using the ‘global’ option from autoreject (Jas et al., 2017). We then averaged the artefact-evoked signal (see ‘average’ option in `mne.preprocessing.compute_proj_ecg`) to enhance subspace estimation and only considered one single projection vector to preserve as much signal as possible.

#### Rejection of residual artifacts

To avoid contamination with artifacts that were not removed by SSS or SSP, we used the ‘global’ option from autoreject (Jas et al., 2017). This yielded a data-driven selection of the amplitude range above which data segments were excluded from the analysis.

#### Temporal Filtering

To study band-limited brain dynamics, we applied bandpass-filtering using the frequency band definitions in Table 2. We used default filter settings from the MNE software (development version 0.19) with a windowed time-domain design (firwin) and Hamming taper. Filter length and transition band-width was set using the ‘auto’ option and depended on the data.

#### Epoching

For the active and passive tasks, we considered time windows between −200 to 700 milliseconds around stimulus-onset and decimated the signal by retaining every eighth time sample. Baseline correction was applied based on the time window between −200 to 0 milliseconds. For resting-state, we considered sliding windows of 5 seconds duration with no overlap and no baseline correction. To reduce computation time, we retained the first 5 minutes of the recording and decimated the signal by retaining every fifth time sample.

#### Channel selection

It is important to highlight that after SSS, the magnetometer and gradiometer data are reprojected from a common lower dimensional SSS coordinate system that typically spans between 64 and 80 dimensions. As a result, both sensor types contain highly similar information, which also modifies the inter-channel correlation structure (Garcés et al., 2017). The MNE software, by default, treats them as a single sensor type in many of the analyses that follow and uses as degrees of freedom the number of underlying SSS dimensions. To simplify computation, we constrained the analysis to magnetometers. For some aspects of feature engineering in sensor space, i.e., extraction of peak latency, we used the gradiometers as they tend to yield a cleaner view on the signal.

#### Covariance Modeling

To control the risk of overfitting in covariance modeling (Engemann and Gramfort, 2015), we used a penalized maximum-likelihood estimator implementing James-Stein shrinkage (James and Stein, 1992) of the form

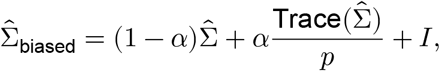

where *α* is the regularization strength, 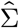 is the unbiased maximum-likelihood estimator and *p* is the number of features. This, intuitively, amounts to pushing the covariance towards the identity matrix. Here, we used the Oracle Approximation Shrinkage (OAS) (Chen et al., 2010) to compute the shrinkage factor *α* mathematically.

#### Source Localization

To estimate cortical generators of the MEG signal, we employed the cortically constraint Minimum-Norm-Estimates (Hämäläinen and Ilmoniemi, 1994) based on individual anatomy of the subjects. If no additional preprocessing is applied, the resulting projection operator depends exclusively on the anatomy of the subject and can be expressed as

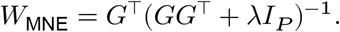

Here 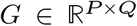 denotes the forward model quantifying the spread from sources to M/EEG observations and λ a regularization parameter that controls the spatial complexity of the model. The forward model is obtained by numerically solving Maxwell’s equations based on the estimated head geometry, which we obtained from the Freesurfer brain segmentation. We estimated the source amplitudes on a grid of 8,196 equally spaced candidate dipole locations. We used spatial whitening to approximate the model assumption of Gaussian noise. The whitening operator was based on the empty room noise covariance and applied to the MEG signal and the forward model. We applied no noise normalization and used the default depth weighting (Lin et al., 2006) as implemented in the MNE software (Gramfort et al., 2014) with weighting factor of 0.8 (Lin et al., 2006) and a loose-constraint of 0.2. The regularization parameter λ^2^ was expressed with regard to the signal-to-noise ratio and kept at the default value of 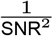 with SNR = 3.

## MRI data processing

### Data acquisition

For additional details on data acquisition, please consider the reference publications of the CAM-Can (Taylor et al., 2017; Shafto et al., 2014). The following sections will describe the custom data processing conducted in our study.

### structural MRI

For preprocessing of structural MRI data we used the FreeSurfer software (http://surfer.nmr.mgh.harvard.edu/)) (Fischl, 2012). Reconstruction included the following steps (adapted from the methods citation recommended by the authors of FreeSurfer): motion correction and average of multiple volumetric T1-weighted images (Reuter et al., 2010), removal of non-brain tissue (Segonne et al., 2004), automated Talairach transformation, segmentation of the subcortical white matter and deep gray matter volumetric structures (Fischl et al., 2002, 2004) intensity normalization (Sled et al., 1998), tessellation of the gray-matter / white-matter boundary, automated topology correction (Fischl et al., 2001; Segonne et al., 2004), and surface deformation following intensity gradients (Dale et al., 1999; Fischl and Dale, 2000). Once cortical were computed, so-called deformable procedures were applied including surface inflation (Fischl et al., 1999), registration to a spherical atlas (Fischl et al., 1999) and cortical parcellation (Desikan et al., 2006).

### fMRI

The available fMRI data were visually inspected. The volumes were excluded from the study provided they had severe imaging artifacts or head movements with amplitude larger than 2 mm. After the rejection of corrupted data we obtained a subset of 626 subjects for further investigation. The fMRI volumes underwent slice timing correction and motion correction to the mean volume. Following that, co-registration between anatomical and function volumes was done for every subject. Finally, brain tissue segmentation was done for every volume and the output data were morphed to the MNI space.

### Scientific Computation and Software

#### Computing environment

For preprocessing and feature-extraction of MEG, MRI and fMRI we used a high-performance Linux server (72 cores, 376GB RAM) running Ubuntu Linux 18.04.1 LTS. For subsequent statistical modeling, we used a golden Apple Mac-Book 12’ (early 2016) running MacOS Mojave (8GB RAM). General purpose computation was carried out using the Python (3.7.3) language and the scientific Python stack including NumPy, SciPy, Pandas, and Matplotlib. For embarrassingly parallel processing we used the joblib library.

#### MEG processing

For MEG processing, we used the MNE-Python software (Gramfort et al., 2014, 2013) (version 0.19 dev). All custom analysis code was scripted in Python and is shared in a dedicated repository including a small library and scripts (see section Code Availability).

#### MRI & fMRI processing

For anatomical reconstruction we used the shell-script based FreeSurfer software Fischl et al. (2002). We used the pypreprocess package, which reimplements parts of the SPM12 software for the analysis of brain images (The Wellcome Centre for Human Neuroimaging, 2018), complemented by the Python-Matlab interface from Nipype (Gorgolewski et al., 2011). For feature extraction and processing related to predictive modeling with MRI and fMRI, we used the NiLearn package (Abraham et al., 2014).

#### Statistical modeling

For predictive modeling, we used the scikit-learn package (Pedregosa et al., 2011) (version 0.21). We used the R (3.5.3) language and its graphical ecosystem (R Core Team, 2019; Wickham, 2016; Slowikowski, 2019; Clarke and Sherrill-Mix, 2017; Canty and Ripley, 2017) for statistical visualization of data.

#### Code Availability

We share all code used for this publication. The code resources for different components can be freely accessed on GitHub in two repositories, one for data processing, feature extraction and predictive modeling^1^, one for statistical analysis and visualization ^2^. Our stacked model architecture can be compactly expressed using the StackingRegressor class in scikit-learn (Pedregosa et al., 2011) as of version 0.22.

## Author contributions

D.A.E., A.G. and G.V. conceived the project. D.A.E. and O.K. conducted the data analysis. G.L. and D.S. contributed analytical solutions. D.A.E. completed the first manuscript draft. All authors provided feedback and revised the manuscript.

## Acknowledgments

This work was partly supported by a 2018 ‘médecine numérique’ (for digital medicine) thesis grant issued by Inserm (French national institute of health and medical research) and Inria (French national research institute for the digital sciences). It was also partly supported by the European Research Council Starting Grant SLAB ERC-YStG-676943.

We thank Sheraz Khan for help with the Freesurfer segmentation and data management of the Cam-CAN dataset. We thank Mehdi Rahim for advice with the model stacking framework and data management of the Cam-CAN dataset. We thank Donald Krieger and Timothy Bardouille for help with the MEG co-registration. We thank Danilo Bzdok for feedback on the first version of the preprint.

## Supporting Information

### Figure 2 supplement

**Fig. 2 – supplement 1.**
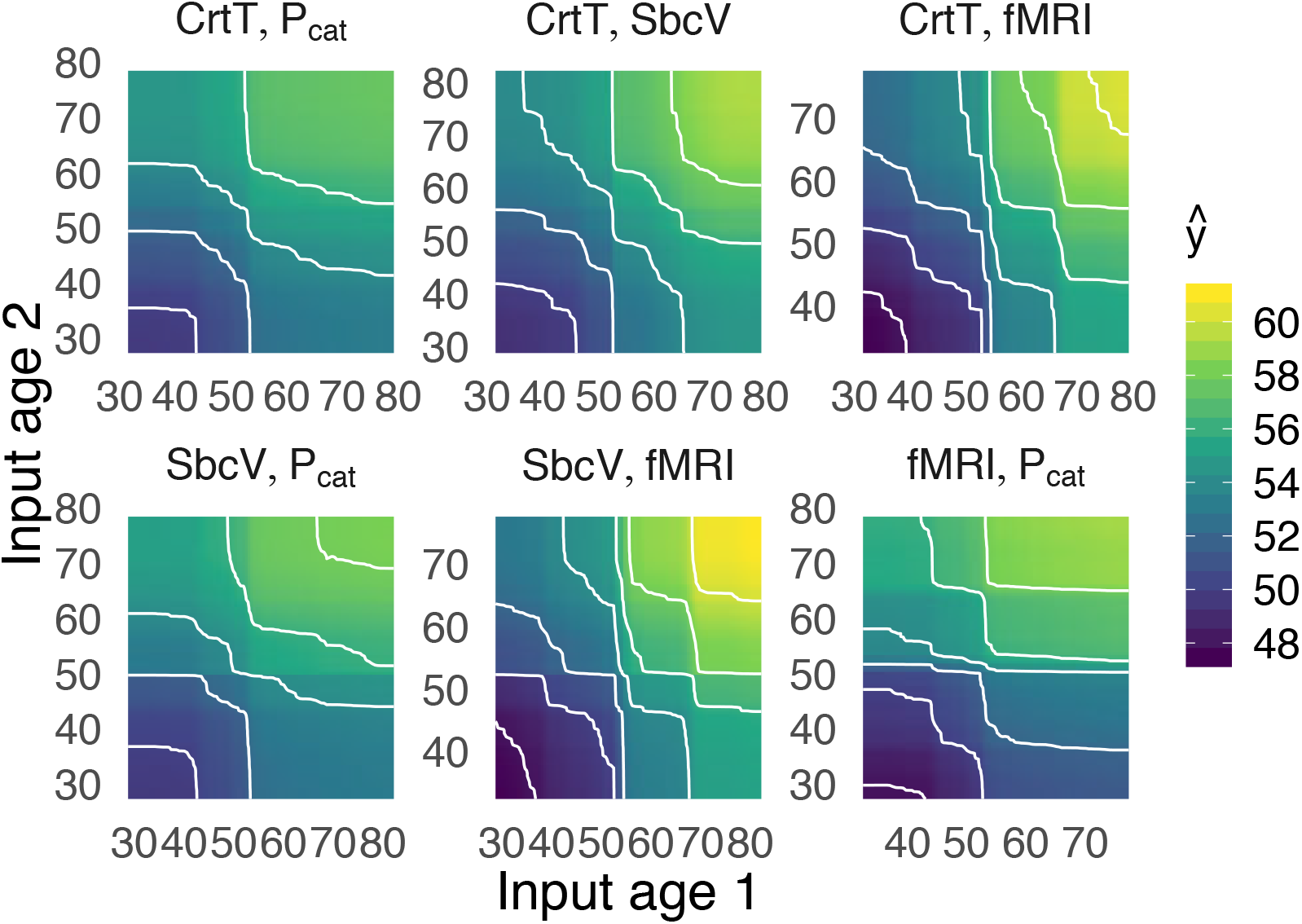
Two-dimensional partial-dependency analysis for top-important stacking inputs. The x and y axes depict the empirical value range of the age inputs. The color and contours show the resulting output prediction of the stacking model. Additive patterns dominating, suggesting independent contributions of MEG and fMRI with little evidence for interaction effects.

**Fig. 2 – supplement 2.**
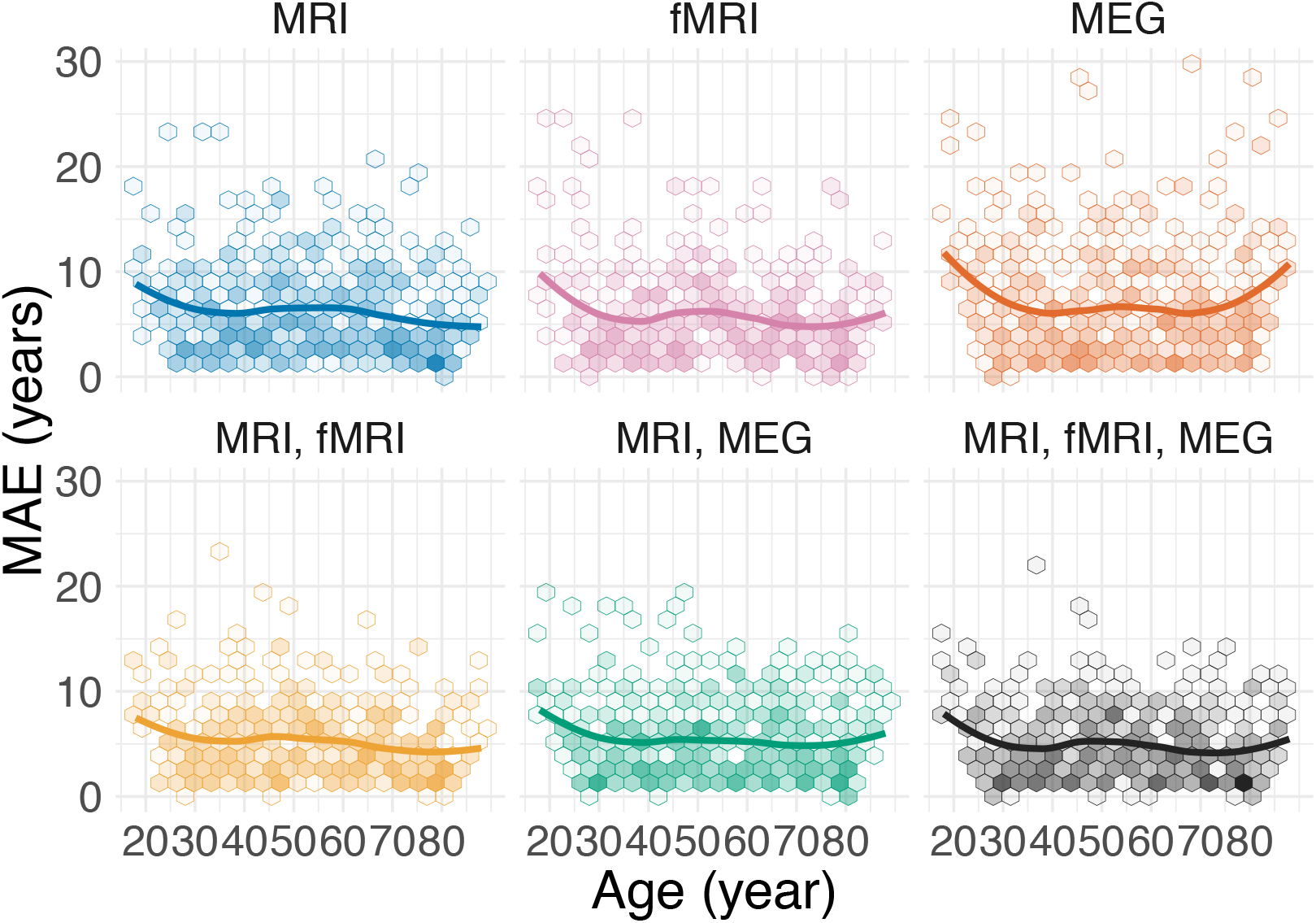
Breakdown of prediction error across age by stacking model. The upper row shows unimodal models, the lower row multimodal ones. Extreme error, especially in young and old subjects was mitigated by stacking.

### Figure 3 supplement

**Fig. 3 – supplement 1.**
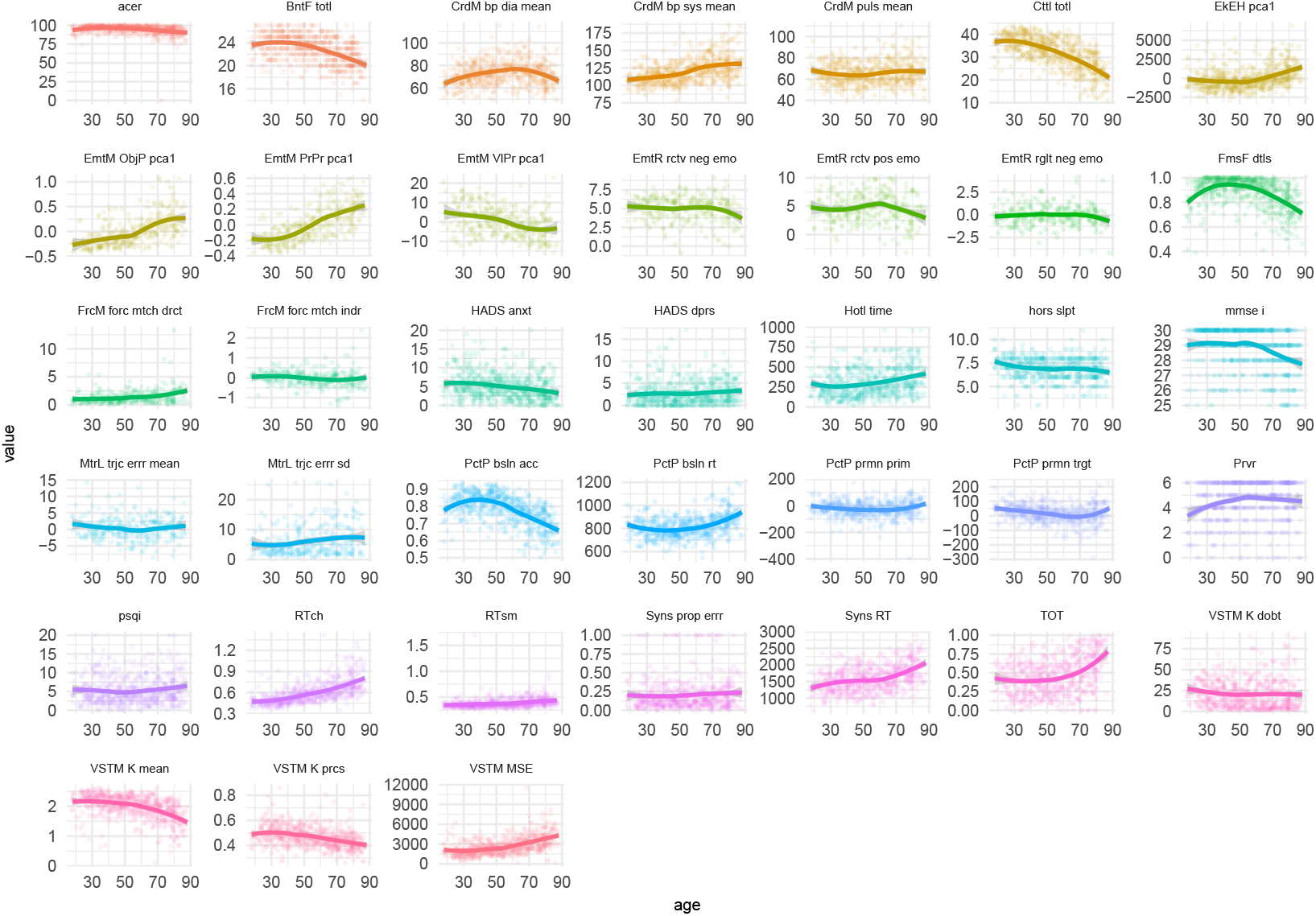
Coefficients of deconfounded linear models predicting neuropsychological scores from brain ageΔ

**Fig. 3 – supplement 2.**
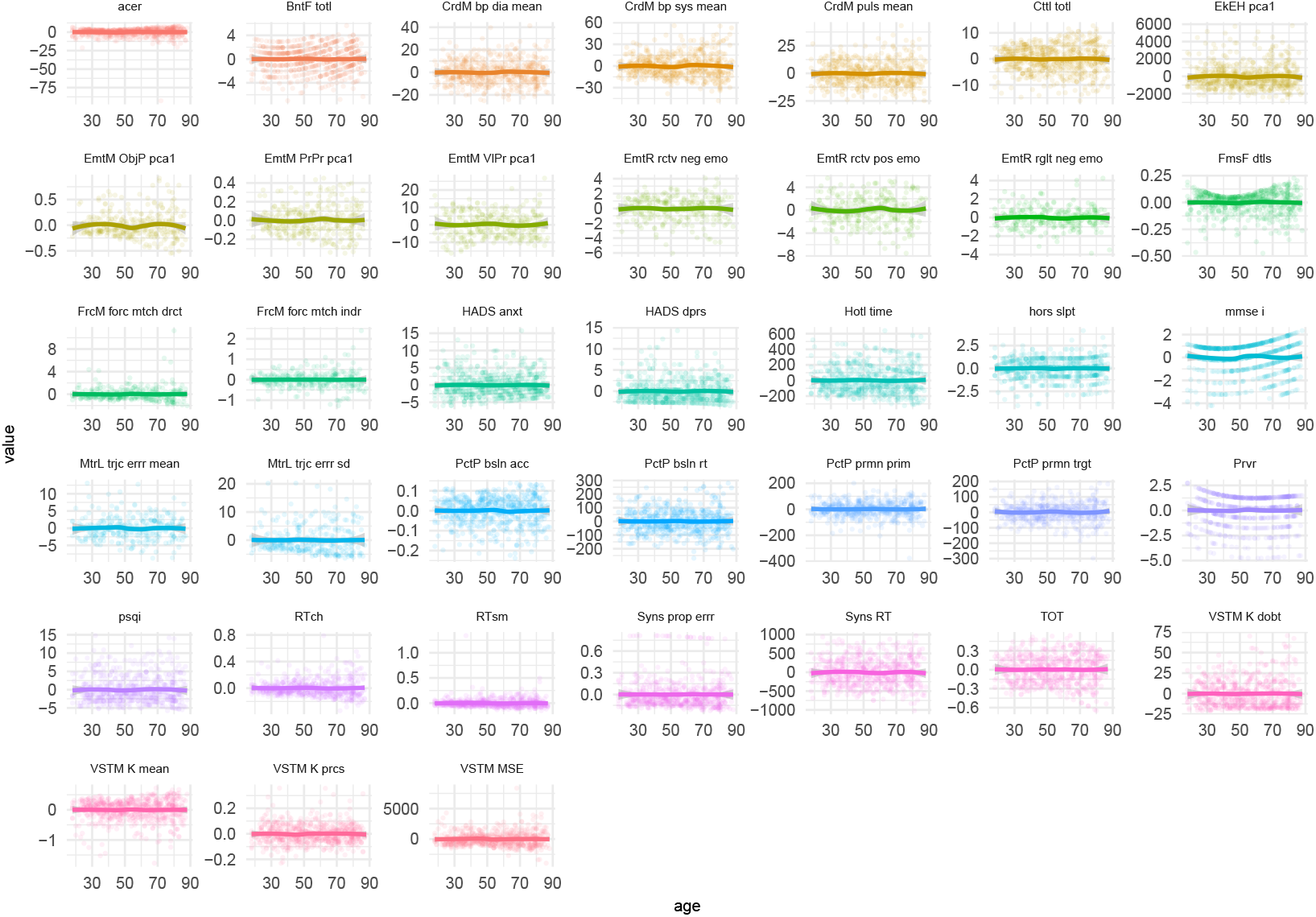
Neuropsychological scores across lifespan after residualizing for age.

**Fig. 3 – supplement 3.**
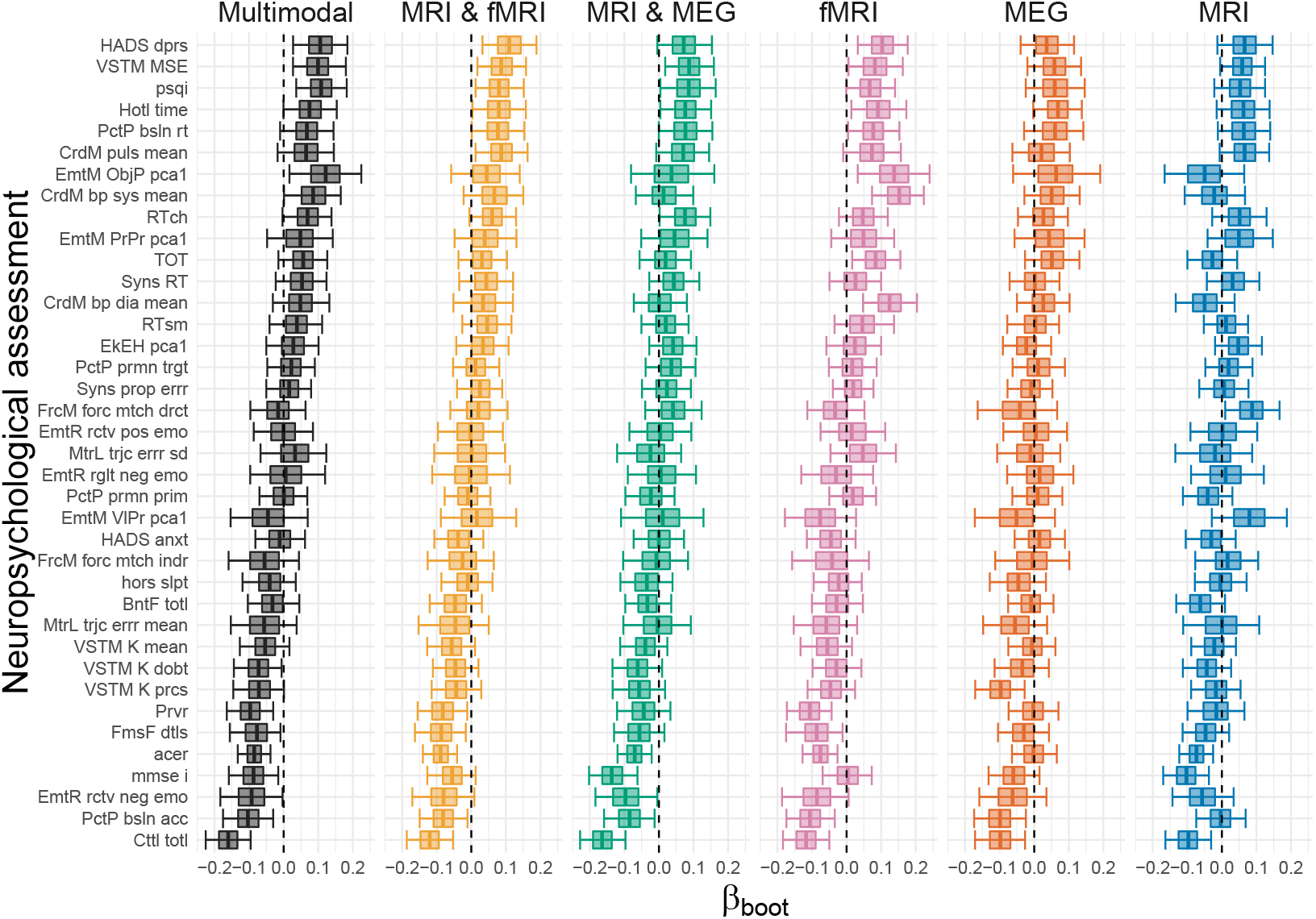
Residual correlation between brain age Δ and neuropsycholgical assessment. The x-axis depicts the coefficients from univariate regression models. Uncertainty estimates are obtained from non-parametric bootstrap estimates with iterations.

### Figure 4 supplement

**Fig. 4 – supplement 1.**
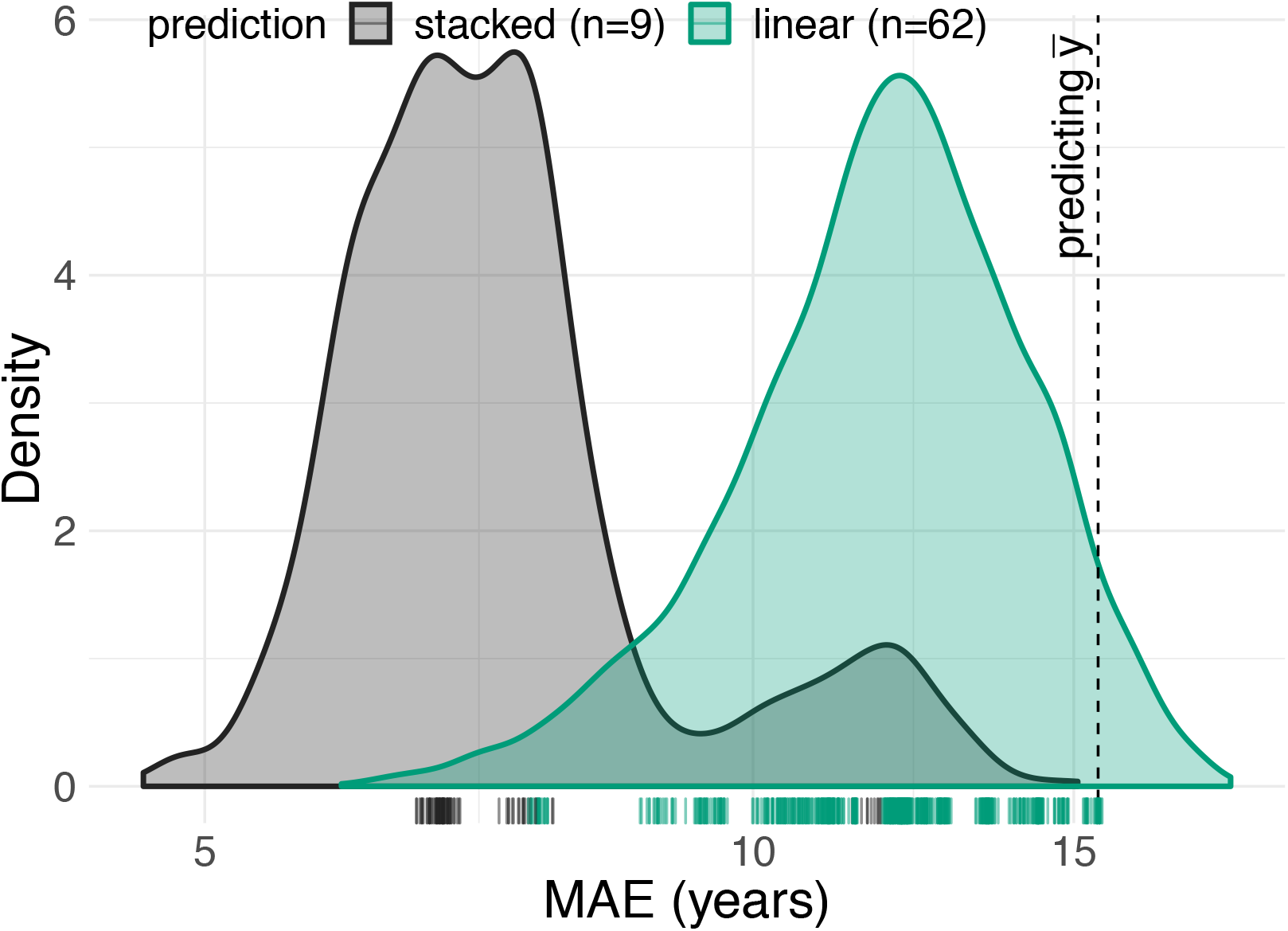
Distribution of prediction errors across 62 first-level linear models (green) and 9 second-level stacking models (black) based on random forests. One can see that stacking mitigates prediction error.

**Fig. 4 – supplement 2.**
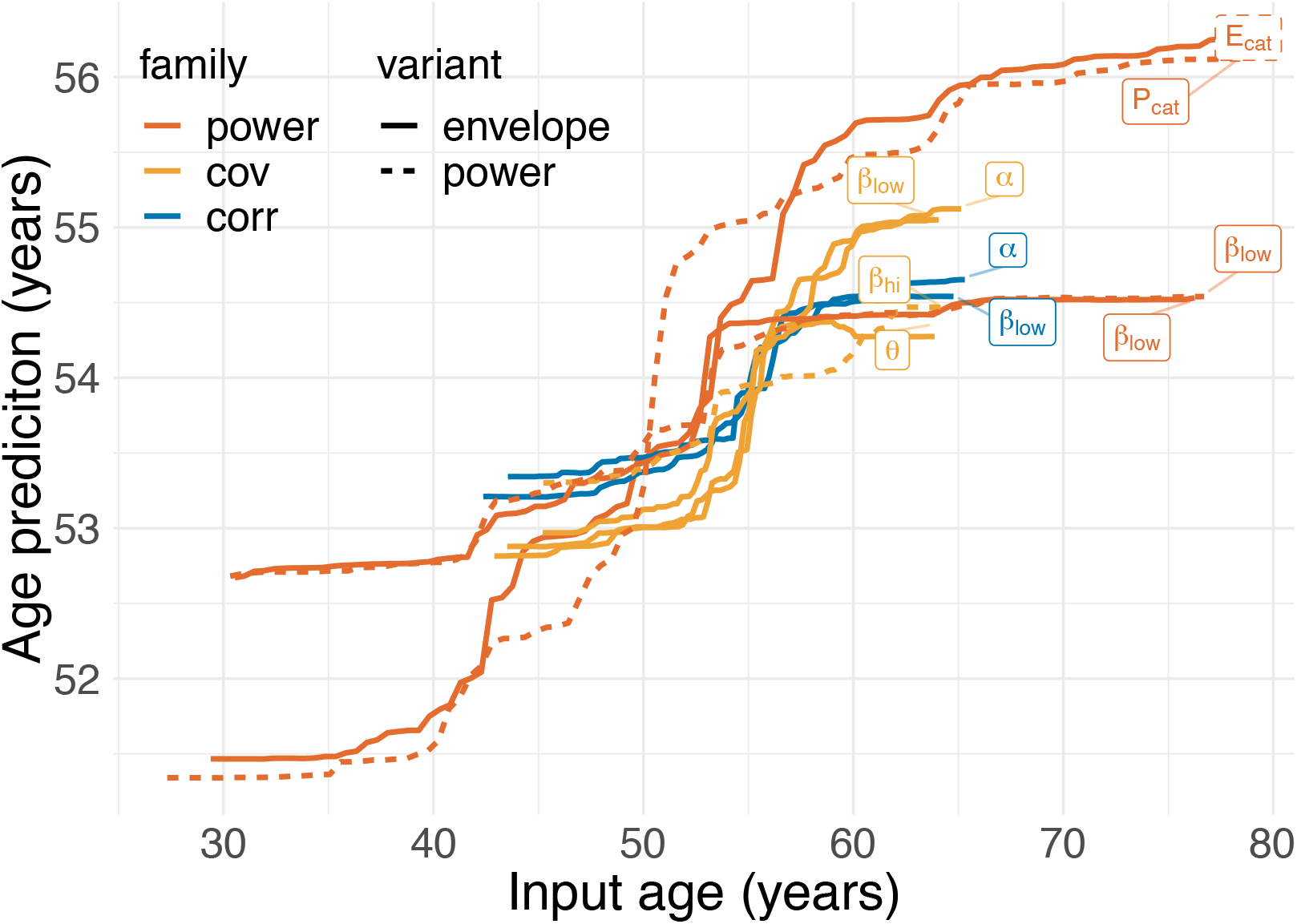
Partial dependence between top age-inputs and the final stacked age-prediction. One can see that extreme input-predictions are pulled toward the mean, following a non-linear step-pattern.

## Footnotes

1 https://github.com/OlehKSS/camcan_analysis

2 https://github.com/dengemann/paper-multimodal-stacking-figures/

